# Involvement of ASIC1a channels in the spinal processing of pain information by deep projection neurons

**DOI:** 10.1101/2021.08.02.454740

**Authors:** Magda Chafaï, Ariane Delrocq, Perrine Inquimbert, Ludivine Pidoux, Kevin Delanoe, Eric Lingueglia, Romain Veltz, Emmanuel Deval

## Abstract

Dorsal horn of the spinal cord is an important crossroad of pain neuraxis, especially for the neuronal plasticity mechanisms that can lead to chronic pain states. Windup is a well-known spinal pain facilitation process initially described several decades ago, but which exact mechanism is still not fully understood. Here, we combine both *ex vivo* and *in vivo* electrophysiological recordings of spinal neurons with computational modelling to demonstrate a role for ASIC1a-containing channels in the windup process. Spinal application of the ASIC1a inhibitory venom peptides mambalgin-1 and psalmotoxin-1 (PcTx1) significantly reduces the ability of deep wide dynamic range (WDR) neurons to develop windup *in vivo*. All deep WDR-like neurons recorded from spinal slices exhibit an ASIC current with biophysical and pharmacological characteristics consistent with functional expression of ASIC1a/ASIC2 heteromeric channels. A computational model of WDR neuron supplemented with heteromeric ASIC1a/ASIC2 channel parameters accurately reproduces the experimental data, further supporting a positive contribution of these channels to windup. It also predicts a calcium-dependent windup decrease for elevated ASIC conductances, a phenomenon that was experimentally validated using either a combination of calcium-activated potassium channel inhibitory peptides (apamin and iberiotoxin), or the Texas coral snake ASIC-activating toxin (MitTx). This study demonstrates a possible dual contribution to windup of calcium permeable ASIC1a/ASIC2 channels in deep laminae projecting neurons, promoting it upon moderate channel activity, but ultimately leading to calcium-dependent windup inhibition associated to potassium channels when activity increases.

## Introduction

Dorsal horn of the spinal cord is a key point of the pain neuraxis where sensory-nociceptive information, coming from the periphery, enters the central nervous system to be integrated, processed, and sent to the brain. It consists of an extremely complex neuronal network organized in different laminae (laminae I to VI), including different types of projection neurons as well as excitatory/inhibitory interneurons (for review, see (1)). The complexity of dorsal spinal cord neuronal network not only lies on its architecture and its great neuronal diversity, but also on the fact it receives various sensory and nociceptive inputs from the periphery, and that it is modulated by descending pathways from supra-spinal levels. Spinal inputs come from peripheral Aβ, Aδ and C fibers and, importantly, these inputs can be subject to different facilitation/sensitization processes, leading to pain hypersensitivity and allodynia (for reviews, see (2, 3)). These processes generally result from intense and repetitive noxious inputs and are associated with neuronal plasticity that sensitizes spinal neurons by developing/increasing their spontaneous activity, decreasing their activation threshold, amplifying their response to stimuli and/or enlarging their receptive fields. Sensitized states can be long-lasting but are normally not permanent. However, they can also be associated to chronic pain states of clinical relevance (4), when pain loses its physiological protective function to fall into pathology.

Many progresses have been made over the last decades in the understanding of spinal facilitation/sensitization molecular mechanisms, including windup, which is a “short term” facilitation process typical of wide dynamic range (WDR) projecting neurons (5). Windup is a homosynaptic facilitation process of C-fiber inputs following peripheral low-frequency repetitive stimulations, resulting in a progressive increase of the number of action potentials (APs) evoked by WDR neurons (2). Although windup and central sensitization share common properties, they are not equivalent but windup can lead to some aspects of central sensitization (6). Therefore, windup remains an interesting way to study the processing of nociceptive information by spinal cord neurons (for reviews see, (2, 7)). Many factors have been reported to contribute to and/or to modulate windup, among which the most important appear to be NMDA receptors (8, 9), NK1 receptors (10) and L-type calcium channels (11-13).

ASICs (Acid-Sensing Ion channels) are voltage-independent cationic channels mainly selective for Na^+^ ions (14). These channels are gated by protons, *i*.*e*., they are sensors of extracellular pH. Several ASIC subunits have been identified in mammals (ASIC1 to ASIC4, for reviews see (15, 16)), which assemble as trimers (17) to form functional channels, including homomers and heteromers (18). ASICs are widely expressed in the nervous system and are found throughout the pain neuraxis, including spinal neurons of the dorsal horn (19-21). The discovery and *in vivo* use of pharmacological tools able to modulate their activities (for review see (22)) have largely contributed to fuel the body of proofs arguing for the involvement of these channels in pain, both in human (23, 24), and mainly in animal models of pain, either at the peripheral or the central level (25-30). Interestingly, peptides isolated from animal venoms, which remains so far the most specific pharmacological tools able to modulate particular subtypes of ASICs (28, 31-33), were reported to have strong analgesic or painful effects in animals (25-28, 33-35), depending on whether they inhibit or activate the channels, respectively. ASICs are thus interesting pharmacological targets for pain, but if their role in peripheral sensory neurons is relatively well documented, little is known about their participation to pain processes in the central nervous system. For instance, while pharmacological inhibition of ASICs at the spinal cord level is known to produces potent analgesia (20, 28, 34), the mechanism of this effect still remains poorly understood.

In this work, we investigated the role of spinal ASICs, which are mainly formed by ASIC1a-containing channels (19-21), in the processing of pain information. By combining both *in vivo* and *ex vivo* neurophysiological approaches with pharmacology and computational modeling, we demonstrate that ASIC1a heteromeric channels in deep laminae projecting neurons participate to the spinal windup facilitation process.

## RESULTS

### Pharmacological inhibition of ASIC1a-type channels in the dorsal spinal cord decreases the evoked activity of projection neurons

To study the *in vivo* role of spinal ASICs in the processing of sensory and nociceptive inputs, extracellular recordings of dorsal horn neurons (DH neurons) were performed on anesthetized rats (Fig. 1 and supplementary Figs. 1-2). ASIC1a homomeric and ASIC1a/ASIC2a heteromeric channels have been reported to be predominant in dorsal spinal cord neurons (19-21). *In vivo* recordings of DH neurons were thus combined with the use of mambalgin-1 and PcTx1, two inhibitory peptides specific for ASIC1-type channels, including ASIC1a homomers (28, 31) and ASIC1a/ASIC2 heteromers (28, 36-38), with nanomolar to few hundred nanomolar affinities. The combination of two independent selective inhibitors both increased the specificity of our results and allowed to gain information on the ASIC1a channel subtypes involved because of the overlapping pharmacological profiles of the two inhibitors. WDR neurons were classically identified according to their ability to evoke action potential (AP) firing in response to both innocuous (brush) and noxious (pinch) mechanical stimulations of their receptive fields on rat hind-paws (Fig. 1A). A windup protocol was designed (16 repetitive electrical stimulations at 1 Hz) and applied onto the receptive fields of WDR neurons every 10 minutes, before (control) and after spinal application of mambalgin-1 (Fig. 1B-E) or PcTx1 (Fig. 1F-H). In control conditions, the number of APs evoked by C-fibers increased, as expected, with the number of stimulation, and the maximal windup was reached between the 13^th^ and the 16^th^ stimulation (Figs. 1C and 1F). Spinal application of mambalgin-1 for 10 min drastically reduced the C-fiber induced windup by approximately 50% (Fig. 1C), *i*.*e*., 44% and 55% depending on whether the percentage of inhibition was calculated from the total number of AP (Fig. 1D), or from the area under curve (Fig. 1E), respectively. Interestingly, it also slightly but significantly decreased by 21% the number of APs of the C-components at the input, which corresponds to the number of APs evoked by WDR neurons at the first stimulation (Fig. 1C, inset). The same kind of effects was also obtained when PcTx1 was delivered at the spinal level (Fig 1F-H). Indeed, windup was inhibited by 57% or 60% in terms of total number of AP or AUC, respectively (Fig. 1G and H). Moreover, PcTx1 also affected the input, with a 47% decrease of the number of AP induced at the first stimulation (Fig. 1F, *inset*).

**Figure 1:**
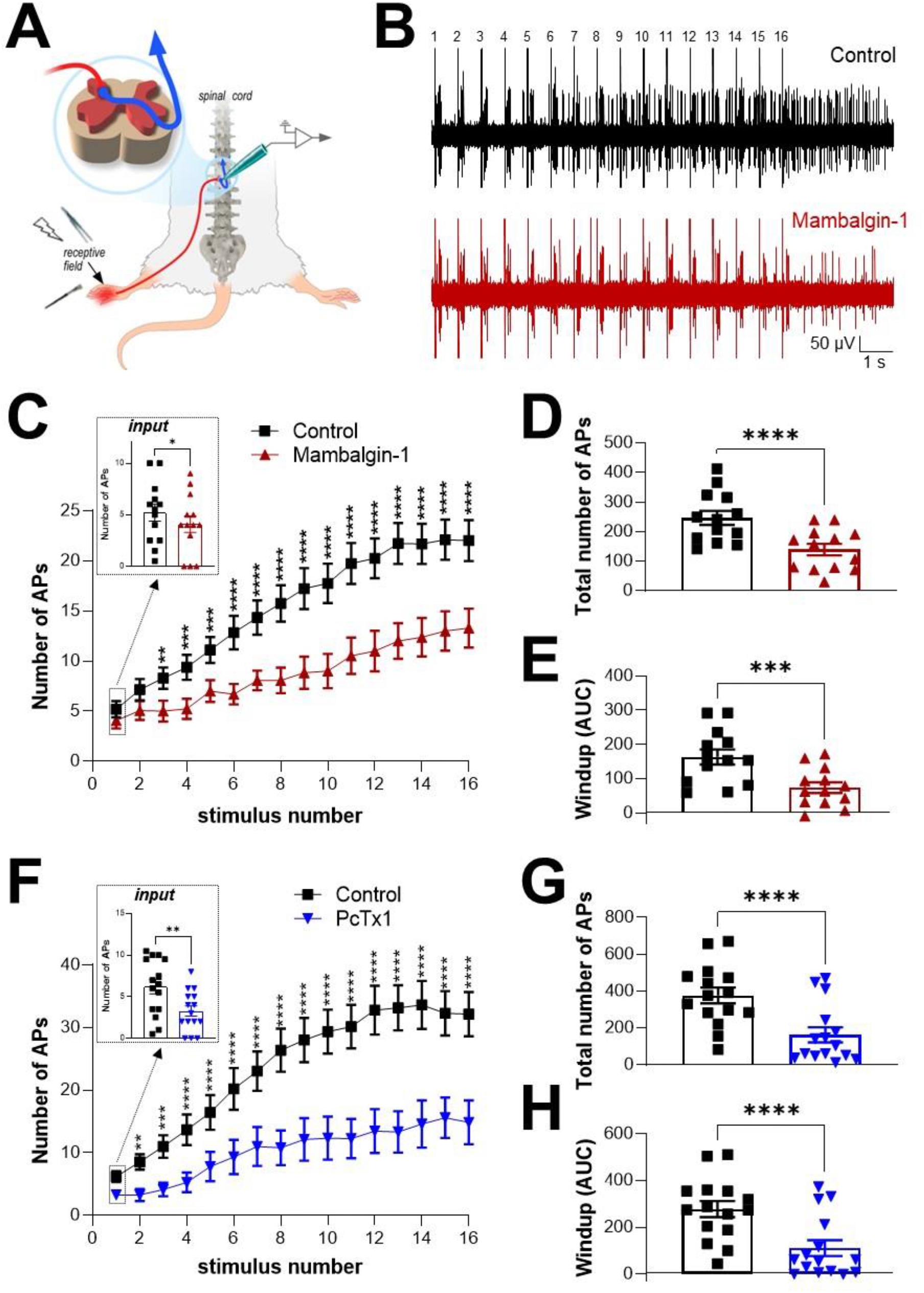
Windup inhibition induced by spinal application of ASIC1a channel blockers. ***A***, Schematic representation of the *in vivo* electrophysiological method used to record WDR neurons from anesthetized rat. Receptive fields of neurons are stimulated either mechanically (brush, pinch) or electrically. ***B***, Typical electrophysiological recording of a WDR neuron obtained following 16 electrical repetitive stimulations of its receptive field at 1Hz (windup protocol, see methods), before (control, top trace) and after spinal application of 30µM mambalgin-1 (bottom trace). ***C***, Windup curves representing the number of C fiber-evoked APs for each of the 16 repetitive electrical stimulations in basal conditions (control), and 10 min after spinal application of mambalgin-1 (n=13, **, *** and **** significantly different with *p*<0.01, *p*<0.001 and *p*<0.0001, respectively, two-way ANOVA followed by a Sidak multiple comparison test). The *inset* shows the average number of APs evoked by the first electrical stimulation, *i*.*e*., before establishment of windup (n=13, \**p*<0.05, paired t test). ***D and E***, Statistical analyses of the total number of AP evoked by C fibers during a windup protocol (D), and of the global windup estimated as the area under curves (AUC, E) from the data shown C (n=13, \*\*\**p*<0.001 and \*\*\*\**p*<0.0001, paired t tests). ***F***, Windup curves representing the number of C fiber-evoked AP for each of the 16 repetitive electrical stimulations in control condition and after spinal application of 30µM PcTx1 (n=15, **, *** and **** significantly different with *p*<0.01, *p*<0.001 and *p*<0.0001, respectively, two-way ANOVA followed by a Sidak multiple comparison test). The *inset* shows the average number of AP evoked by the first electrical stimulation (n=15, \*\**p*<0.01, paired t test). ***G and H***, Statistical analyses of the total number of AP evoked by C fibers during a windup protocol (G), and of the global windup estimated as the area under curves (AUC, H) from the data shown F (n=15, \*\*\*\**p*<0.0001, paired t tests).

Spinal application of mambalgin-1 or PcTx1 also affected the Aβ- and Aδ-evoked activities in WDR neurons (Supplementary Fig. 1), with small inhibitions of associated firings. Indeed, the firing induced by brushing (Supplementary Fig. 1A), which can be related to Aβ-fibers, was decrease by 20% in the presence of mambalgin-1, while PcTx1 had no effect (Supplementary Fig. 1B, *p*=0.07 and *p*=0.31, respectively). Moreover, both mambalgin-1 and PcTx1 significantly reduced the total number of AP induced by Aδ-fibers during the windup protocol (by 26% and 44%, respectively, Supplementary Fig. 1C-D).

All together, these results obtained with two independent selective inhibitors of ASIC1a channel subtypes demonstrate that spinal ASIC1a contribute to the integration of C-fiber inputs and associated windup. In addition, an involvement of either ASIC1a homomers or ASIC1a/ASIC2 heteromers is suggested from the overlapping pharmacological profiles of mambalgin-1 and PcTx1, which strongly affect ASIC1a homomeric channels (28, 31) and have also been reported to potently inhibit ASIC1a/ASIC2 heteromers (36-38).

### ASIC1a channel subtypes are functionally expressed in large neurons from deep laminae V

To further investigate the contribution of spinal ASIC channels to WDR activity, patch-clamp experiments were performed on spinal cord slices (Fig. 2). Electrophysiological recordings were made in deep laminae V to record and characterize native ASIC currents in large neurons, which most likely correspond to the WDR neurons (39) studied *in vivo* (Fig. 1). All the neurons recorded displayed an ASIC-type current in response to extracellular acidification from pH7.4 to pH6.6 (Fig. 2A), with an average amplitude of 242 ± 37pA and an inactivation time constant of 1,536 ± 98ms (Fig. 2B, left panel). Such inactivation kinetics were in the range of those of homomeric ASIC1a (Fig. 2B, right panel) and heteromeric ASIC1a/ASIC2 (18, 26) channels. On the other hand, the native ASIC current had a stable amplitude over time and no tachyphylaxis phenomenon (36, 40) was observed upon repetitive activation at pH6.6 (Fig. 2C), which was different from what has been described for ASIC1a homomeric channels (36, 40), and to what was observed here in HEK293 cells transfected with ASIC1a (Fig. 2C, inset). Tachyphylaxis is absent in ASIC1a heteromeric channels containing the ASIC2 subunit (36, 40), suggesting that the native ASIC currents recorded from deep laminae neurons were carried by ASIC1a/ASIC2 heteromers. Finally, native ASIC currents were almost completely blocked by extracellular applications of PcTx1 (30nM) or mambalgin-1 (1µM) (Fig. 2A and 2D).

**Figure 2:**
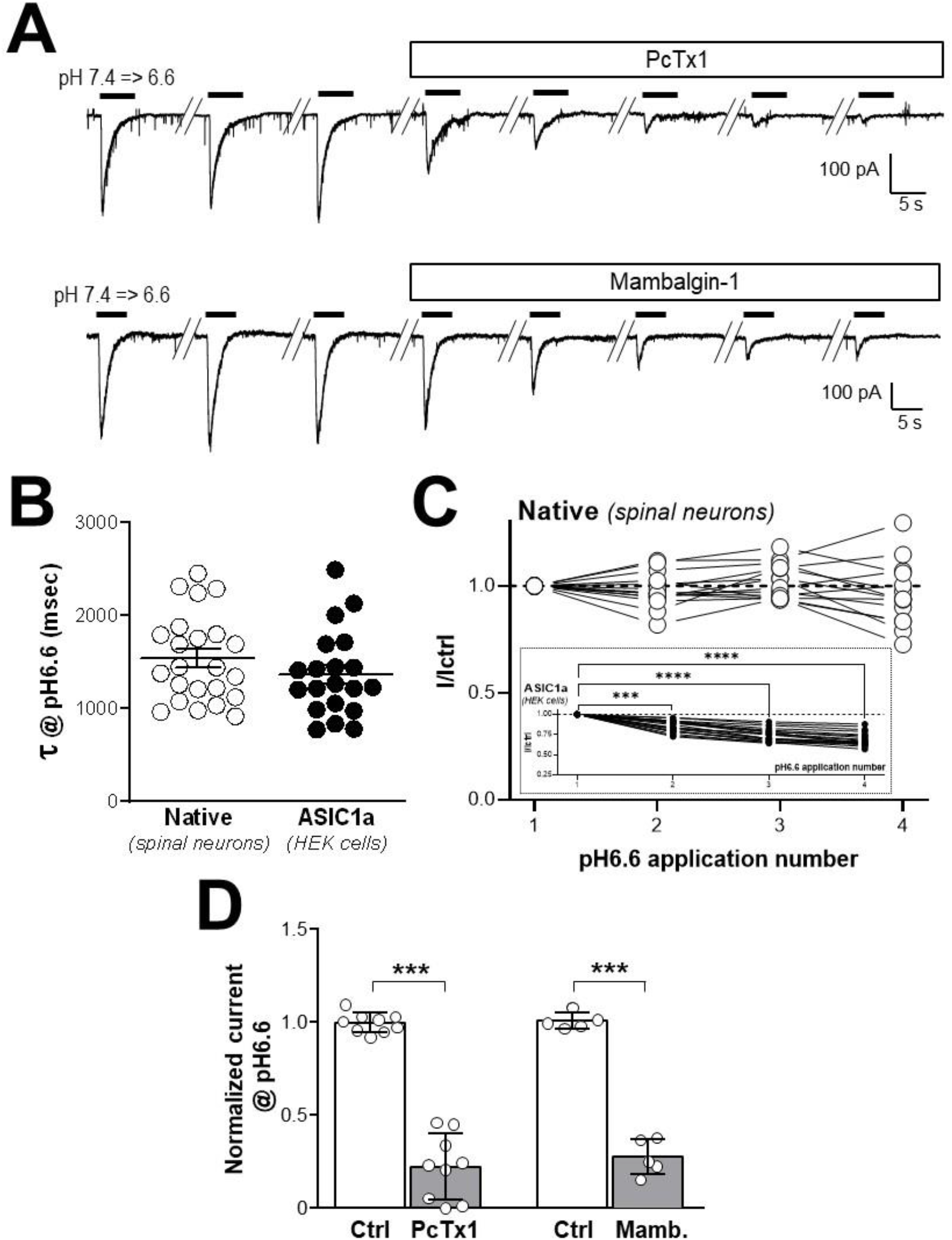
Characterization of a native ASIC1a-type current in deep projection neurons from laminae V. ***A***, Typical voltage-clamp recordings of deep projection neurons (laminae V) obtained from spinal cord slices following extracellular acidifications from pH7.4 to pH6.6. Extracellular applications of the ASIC1a inhibitory peptides PcTx1 (30nM) or mambalgin-1 (1µM) both decrease the amplitude of native pH6.6-induced currents. ***B***, Inactivation kinetics of native pH6.6-induced currents recorded in spinal neurons (native) are compared to those of ASIC1a homomeric currents form HEK293 transfected cells (ASIC1a). Inactivation rates (τ) were obtained by fitting current inactivation decays with a mono-exponential (n=23 and 20 for spinal neurons and ASIC1a-HEK293 cells, respectively, *p*=0.2399, unpaired *t*-test). ***C***, Peak amplitude of the native ASIC current recorded in neurons following four consecutive pH6.6 extracellular acidifications (every 60s). Amplitudes are normalized to the first pH6.6-evoked current (n=14, no significant tachyphylaxis process with *p*=01882, one-way ANOVA test followed by a Dunnet’s *post hoc* test). *Inset* shows the same experiment performed on ASIC1a homomeric current recorded in HEK293 cells (n=20, significant tachyphylaxis process with \*\*\**p*<0.001 and \*\*\*\**p*<0.0001, one-way ANOVA test followed by a Dunnet’s *post hoc* test). ***D***, Statistical analysis of both PcTx1 (30nM) and mambalgin-1 (1µM) effects on the native pH6.6-induced current amplitudes recorded as in A (n=9 and 5, respectively, \*\*\**p*<0.001, paired Student t-test).

Altogether, these data demonstrate for the first time that ASIC1a channel subtypes are functionally expressed in deep projection neurons from laminae V, which is fully consistent with their participation to C-fiber input integration and windup. The properties of native ASIC1a-type currents, *i*.*e*. kinetics, tachyphylaxis and pharmacological sensitivity, suggest the expression of ASIC1a heteromers *(i*.*e*., ASIC1a/ASIC2) rather than homomers.

### Computational model of WDR neuron including ASIC1a channel parameters and acidification of the synaptic cleft

A mathematical model of WDR neurons and windup was initially described by Aguiar and colleagues (41), and was later taken over by Radwani and colleagues (13) to help demonstrate the involvement of Cav1.3 channels in the genesis of windup. We decided to take advantage of this model to help us understand the role of ASIC1a channels in the processing of C-fiber input and windup in WDR neurons (Fig. 3 and Supplementary Fig. 2). In this model, a WDR neuron receives a direct input from an Aδ-fiber as well as an input from a C-fiber through an interneuron (Supplementary Fig. 2A). With this model, we were able to fully reproduce the windup described by Aguiar and colleagues (41), including the windup inhibition when NK1 ion channel parameters were modified (Supplementary Fig. 2B). The C-fiber-induced spiking activity predicted by the WDR neuron model was then compared to our experimental results, *i*.*e*., an average of about 30 neurons recorded in control conditions. The spike time plot (Supplementary Fig. 2C and D) indicated that the model fitted better to experimental data when the interneuron between C-fibers and WDR neuron was removed. We thus decided to use a simplified model without interneuron (Fig. 3A) to further study the involvement of ASIC1a in the windup process.

**Figure 3:**
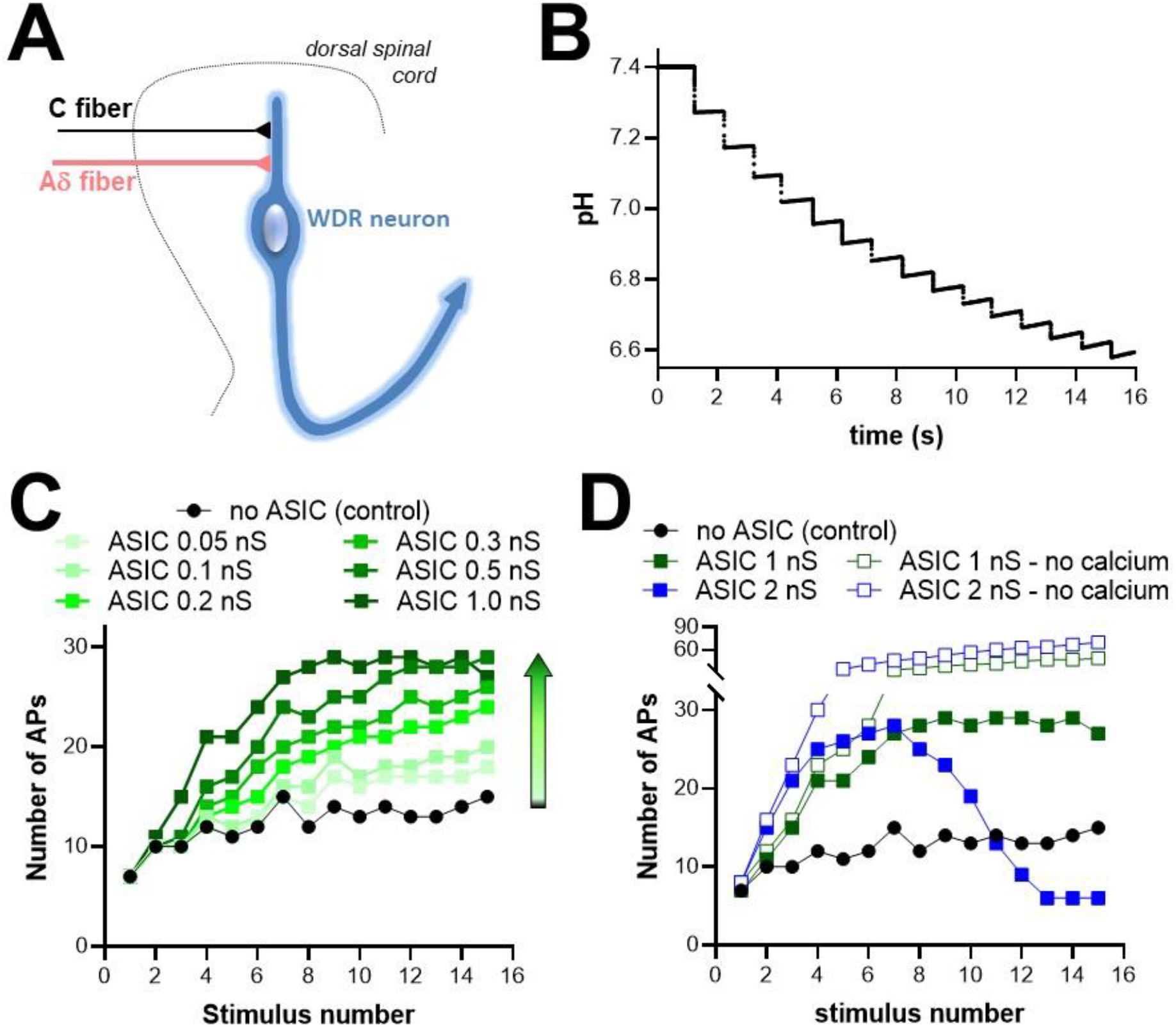
Computational model of WDR neuron with ASIC1a heteromeric channel parameters and a synaptic cleft acidification system. ***A***, Schematic representation of the computational model used, with a WDR projection neuron receiving directly a 20-synapse connection from both Aδ-fibers and C-fibers. ***B***, Simulation curve showing the evolution of synaptic pH as a function of time during windup-inducing stimulations (for the control model, in absence of ASIC channels). ***C***, Simulation windup curves obtained without (control, no ASIC, black dots) and with moderate heteromeric ASIC1a conductances (0.05nS, 0.1nS, 0.2nS, 0.3nS, 0.5nS and 1.0nS, green squares). ***D***, Simulation windup curves obtained in control condition (no ASIC, black dots) and with high heteromeric ASIC1a conductances (1nS, and 2nS, full squares). Compared to control, gradually adding heteromeric ASIC channels first increases windup, then decreases it. Removing the calcium conductance in ASIC parameters (same conductances, open squares) suppresses the inhibitory effect of high ASIC conductances and strongly potentiates windup.

The WDR model was supplemented with either ASIC1a homomeric or ASIC heteromeric/native channel parameters based on a recently described model (42) and previous data (19) (see material and methods). A small calcium conductance was included to account for reported ASIC-mediated intracellular calcium increase. Furthermore, as ASICs are proton-gated ion channels, we also needed to introduce a model of synaptic cleft acidification in our WDR neuron model. This particular point was achieved according to the work of Highstein and colleagues (43), where the synaptic pH is controlled by *(i)* a buffer, *(ii)* a homeostatic system that tends to bring the pH to the physiological value of 7.4 and, *(iii)* a proton current entering the synapse that generates acidification during synaptic activity (see material and methods). Using this model of synaptic pH control, the predicted acidification during a windup protocol is described in Figure 3B.

Including ASIC1a homomeric channel parameters in our WDR neuron model had little or no effect on windup (see methods). In contrast, a windup increase was observed when heteromeric channel parameters were progressively included (up to a 1 nS conductance, Fig. 3C), in good agreement with experimental data showing a windup decrease following spinal application of mambalgin-1 or PcTx1 (Fig. 1). However, when the heteromeric ASIC1a conductance was set at higher levels (above 1nS) the model predicted windup inhibition (Fig. 3D). Interestingly, this effect was dependent on the Ca^2+^ permeability of ASICs since turning this permeability off restored the windup increase (Fig. 3D). This windup inhibition associated with ASIC1a calcium conductance was most likely due to the hyperpolarizing effect of calcium-dependent K^+^ channels present in the model.

The mathematical model described here was thus fully consistent with an involvement in the windup process of heteromeric, rather than homomeric, ASIC1a-containing channels located in WDR neurons at the post-synapse with C-fibers. It also suggested a “bell-shape” participation of ASIC1a to windup, with both positive and negative effects associated to low/medium and high conductances, respectively.

### Maximal activation of spinal ASIC1a channel subtypes inhibits windup through calcium-dependent K^+^ channels

Our mathematical model predicted windup inhibition for high conductances of heteromeric ASIC1a channels in WDR neurons, involving ASIC1a-associated calcium entry and likely subsequent activation of calcium-dependent K^+^ channels. This hypothesis was tested experimentally by using MitTx (Fig. 4), a peptide toxin with potent activating effects on ASICs and particularly ASIC1a channels (33). MitTx was indeed reported to generate a maximal and persistent activation of the channels as well as a potent calcium response in neurons, which was abolished in ASIC1a knockout mice. Spinal application of MitTx induced a dose-dependent inhibition of windup (Fig. 4A and B), in agreement with the effect predicted by our computational model. This effect started at 10^−7^M / 5.10^−7^M and reached a maximum at 10^−6^M. Inhibition of windup by MitTx was total and almost fully reversible upon washout (Fig. 4B). The computational model was next configured to mimic the experimental effect of MitTx, *i*.*e*., with a sustained activation of ASIC channels without inactivation (Fig. 4C). This simulation confirmed that MitTx inhibits windup, in good agreement with experimental data.

**Figure 4:**
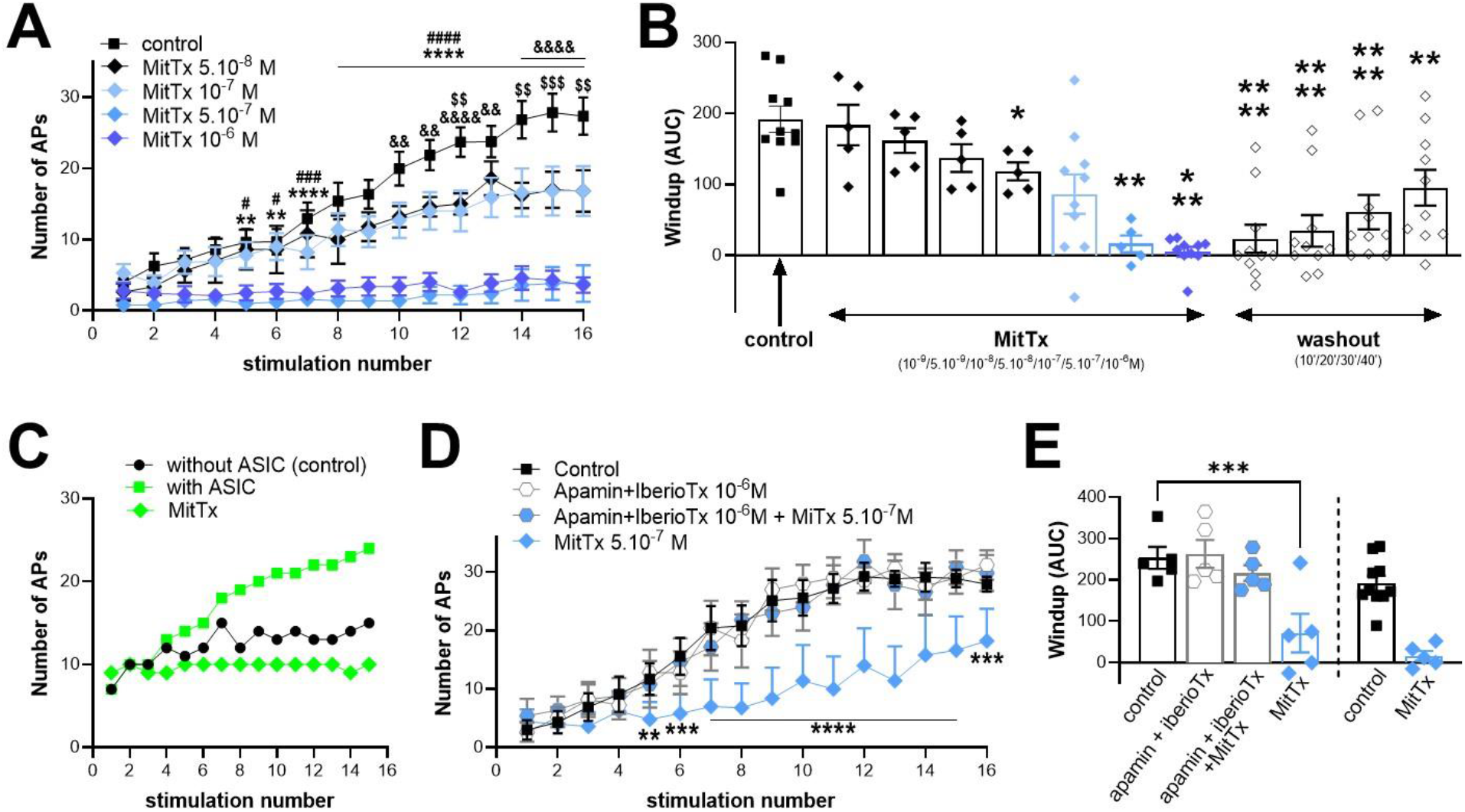
The ASIC1a activator MitTx inhibits windup *in vivo*. ***A-B***, Spinal applications of MitTx dose dependently and reversibly inhibits windup (n=5-10; panel A: \*\**p*<0.01 and \*\*\*\**p*<0.0001 for control *vs*. MitTx 10^−6^ M, ^#^*p*<0.05, ^###^*p*<0.001 and ^####^*p*<0.0001 for control *vs*. MitTx 5.10^−7^ M, ^&&^*p*<0.01 and ^&&&&^*p*<0.0001 for control *vs*. MitTx 10^−7^ M, ^$$^*p*<0.01 and ^$$$^*p*<0.001 for control *vs*. MitTx 5.10^−8^ M; panel B: \**p*<0.05; **p<0.01, p<0.001 and ***p<0.001 as compared to control, respectively; Mixed-effect analyses followed by Dunnet’s multiple comparison tests). ***C***, Simulation of the effect of MitTx on windup (without ASIC: no ASIC conductance; with ASIC: with a heteromeric ASIC conductance of 0.2nS; MitTx: sustained activation of heteromeric ASIC with a same conductance of 0.2nS without inactivation). ***D-E***, The inhibitory effect of MitTx 5.10^−7^M is abolished by spinal pre- and co-application of the KCa blockers apamin and iberiotoxin (D, 10^−6^M each). Removing these two blockers restored the windup inhibition induced by MitTx. As a comparison, data already presented in B and showing the effect of MitTx 5.10-7M applied alone, *i*.*e*., without pre-application of KCa blockers, are also represented on the bargraph in E (n= 5-10; panel D: \*\**p*<0.01, \*\*\**p*<0.001 and ****p<0.0001 as compared to control, respectively, two-way ANOVA followed by a Dunnet’s multiple comparison test; panel E: \*\*\**p*<0.001, one-way ANOVA followed by a Dunnet’s multiple comparison test).

To test the hypothesis that windup inhibition induced by MitTx was due to the opening of calcium-activated K^+^ channels (KCa) following ASIC1a-associated intracellular calcium increase, we inhibited KCa *in vivo* in deep WDR neurons with apamin and iberiotoxin (Fig. 4D). Spinal application of these two toxins together had no significant effect on windup. However co-application of apamin and iberiotoxin together with MitTx prevented the strong windup inhibition induced by MitTx alone (Fig. 4D-E). The removal of KCa blockers restored the potency of MitTx to inhibit windup, with an effect that was comparable to the one initially observed, *i*.*e*., without pre/co-application of the KCa blockers (Fig. 4E).

To summarize, all these data showed that maximal activation of spinal ASIC1a channels by MitTx inhibited windup through a mechanism that is dependent on calcium-activated K^+^ channels. It is in full agreement with the prediction of our mathematical model and further supports both the relevance of the model and the participation of heteromeric ASIC1a/ASIC2 channels to windup in WDR neurons.

## Discussion

The present work describes how ASIC1a channels are involved in the processing of pain message at the level of deep laminae spinal projection neurons. Spinal ASIC1a activity had already been associated to windup and CFA-induced hypersensitivity of dorsal horn neurons (20). However, the channel subtypes as well as the subsets of spinal neurons involved remained to be demonstrated. By combining *in vivo* and *ex vivo* electrophysiological recordings, pharmacology and computational modeling, we propose that ASIC1a/ASIC2 heteromeric channels are functionally expressed in deep projection neurons likely corresponding to WDR neurons, where they are basically involved in the windup process and participate to the post-synaptic integration of nociceptive information coming from the periphery. This is in good agreement with the description of ASIC1a and ASIC2 as the major channel subunits present in spinal cord neurons, and with the predominant functional expression of ASIC1a/ASIC2 heteromers in primary cultures of spinal neurons (19-21).

Post-synaptic expression of ASIC1a channels has been reported in other area of the central nervous system such as nucleus accumbens and amygdala neurons, where the channels have been involved in synaptic transmission (44, 45). However, expression of different ASIC1a channel subtypes, including homomers and heteromers, remains largely unknown, especially in the different subsets of spinal neurons where they are probably finely tuned at the post- and probably also the pre-synaptic levels. We described here that ASIC1a/ASIC2 heteromers are at least post-synaptically expressed in deep WDR neurons to participate to windup through an intriguing “bell shape” mechanism. Indeed, inhibiting or maximally activating ASIC channels both lead to the same result, *i*.*e*., a windup inhibition. The way inhibitors like mambalgin-1 (or PcTx1) and activators like MitTx inhibit windup is different. If the inhibition of ASIC1a channel subtypes (mambalgin-1 or PcTx1 effects) in WDR neurons produced, as expected, a windup decrease most probably associated with decreased depolarization, the consequence of their over-activation (MitTx effect) was unexpected and associated to the calcium conductance of ASIC1a-containing channels. This latter point strongly argues for an expression in WDR neurons of heteromeric ASIC1a/ASIC2b channels, which have a significant Ca^2+^ permeability (38). It can be hypothesized that, *in vivo*, an excessive synaptic acidification related to intense synaptic activity could in some conditions cause an over-activation of ASIC channels, ultimately leading to windup inhibition associated to calcium-sensitive K^+^ channels. This feed-back loop could constitute a protection mechanism that may prevent WDR neuron from over-excitability. Such a protection mechanism could be a more general property of neurons expressing ASIC1a channel subtypes, and especially those having a significant Ca^2+^ conductance like homomeric ASIC1a and heteromeric ASIC1a/ASIC2b channels (14, 38).

The use of computational modeling has been instrumental to the success of our project. Indeed, using one of the very few existing models of WDR neuron and neglecting network effects, we have been able to mathematically reproduce experimentally obtained results, and point towards a biological mechanism realizing predictions, which were further supported by *in vivo* experiments. Such a back and forth exchange between experimental and computational approaches further demonstrates how computational modeling can help understand the complexity of biological events and, in the present case, how unitary mechanisms can be involved in complex physiological processes such as windup. It is noteworthy that this model was able to *(i)* confirm the *in vivo* observations regarding the role of heteromeric ASIC1a channels in windup, and *(ii)* to predict a windup inhibition upon large activation of ASIC1a/ASIC2 channels through calcium-dependent K^+^ channels, which has then been experimentally validated. The key to these achievements was to introduce as few parameters as possible when we modified the model of Aguiar and colleagues (41) by adding a new model of ASIC1a channels, based on recently published data (42), and a synaptic pH model. Indeed, Aguiar underlined in his paper how tedious tuning his model was. Several research axes are now possible, including the numerical dissection of the precise mechanisms by which WDR response shows up and is altered by ASIC1a/ASIC2 channels, which could provide the operating regime of the WDR neurons and point towards simplified/phenomelogical models of these neurons (46, 47). These simplified models of WDR neuron could be embedded in a network and help revisit our assumption regarding the role of the network in shaping the WDR response.

Most if not all spinal neurons have been reported to display a functional ASIC1a channel subtype, including homomers and heteromers made of ASIC1a, ASIC2a and ASIC2b subunits (19-21). Understanding how these different channel subtypes are distributed within the neuronal network of dorsal spinal cord is an essential point to fully demonstrate how ASICs participate to spinal pain mechanisms. This work provides both experimental and computational arguments for an original participation of calcium permeable ASIC1a/ASIC2b channels in WDR neurons to the pain facilitation process of windup. It also set up an ASIC1a-containing WDR computational model that could be very helpful to further explore the role of ASIC channels in the spinal pain network.

## Material and methods

### Animal anesthesia and surgery

Experiments were performed on 5-7 week-old Wistar male rats (Charles River Laboratories) that were housed in a 12 hours light/dark cycle with food and water available *ad libitum*. Experimental procedures used in this study were approved by the Institutional Local Ethical Committee and authorized by the French Ministry of Research according to the European Union regulations (agreement n°02595.01). Animals were anesthetized with isoflurane (Anesteo, France) and placed on a stereotaxic frame (M2E, France) with the head and vertebral column stabilized by ear bars and vertebral clamps, respectively. Limited laminectomies were performed at the T13-L2 segments to expose the dorsal spinal cord and the underlying dura mater was removed.

### *In vivo* recording of dorsal horn neuron activity

Single unit extracellular recordings of lumbar dorsal horn neurons were made using tungsten paralyn-coated electrodes (0.5MΩ impedance, WPI, Europe). The tip of a recording electrode was initially placed at the dorsal surface of the spinal cord using a micromanipulator (M2E, France) and this initial position was set as 0µm on the micromanipulator’s micrometer. The electrode was then progressively moved down into the dorsal horn until the receptive field of a spinal neuron was localized on the ipsilateral plantar hindpaw using mechanical stimulations. Two types of mechanical stimulations were used to characterize spinal projecting neurons, including non-noxious brushing and noxious pinching, and we focused on WDR neurons responding to both brush and pinch stimuli. The depth of the WDR neurons selected for this study was >400µm, most probably corresponding to deep neurons from laminae V. Activities of WDR neurons were sampled at 20 kHz, band-pass filtered (0.3-30 kHz) and amplified using a DAM80 amplifier (WPI, Europe), digitized with a A/D-D/A converter (1401 data acquisition system, Cambridge Electronic Design, Cambridge, UK), and recorded on a computer using Spike 2 software (Cambridge Electronic Design, Cambridge, UK).

### Windup protocol and analysis

Once a WDR neuron was isolated, its receptive field was stimulated every 10 minutes with the following protocol: 10 times brushing, to generate Aβ-evoked response, followed by a train of 16 supraliminar electrical repetitive stimulations (1Hz, 4ms pulse width, Dagan S900 stimulator, Mineapolis, USA) to induce windup. Intensity of currents injected for the windup protocols was determined as the intensity required to evoke less than 10 action potentials (APs) at the first stimulation, corresponding to 1.2-3 times the AP thresholds. Drugs were applied directly to the dorsal surface of the spinal cord bathed in 40µl of ACSF saline solution that contained (in mM): NaCl 119, KCl 2.5, NaH_2_PO_4_ 1.25, MgSO_4_ 1.3, CaCl_2_ 2.5, NaHCO_3_ 26, glucose 11 and HEPES 10 (pH adjusted to 7.4 with NaOH). APs evoked by WDR were classified according to the time frame at which they were emitted following the electrical stimulation artifact, *i*.*e*., 0-20, 20-90 and 90-300 ms for Aβ-, Aδ- and C-inputs, respectively (48). The remaining APs evoked 300-1,000 ms after the stimulation artifact were classified as the post-discharge activity of WDR neurons. Windup calculations were established by counting the number of APs evoked during the C- and post-discharge parts of recordings. Area under curves (AUC) for windup curves were determined using Prism software.

### Patch-clamp experiments on spinal cord slices

Transverse spinal slices (400µm thick) were prepared from male rats (15-28 days old) as described previously (49). Rats were deeply anesthetized with urethane (1.9 g/kg, i.p.) and killed by decapitation. The spinal cord was removed by hydraulic extrusion and washed in ice-cold (≤4°C) sucrose–artificial CSF (ACSF) containing the following (in mM): 248 sucrose, 11 glucose, 26 NaHCO_3_, 2 KCl, 1.25 KH_2_PO_4_, 2 CaCl_2_, 1.3 MgSO_4_ (bubbled with 95% O_2_ and 5% CO_2_). The lumbar segment was embedded in 5% agarose, and 400-μm-thick transverse slices were cut with a vibratome (VT1200S; Leica, Germany). Slices were stored at room temperature in a chamber filled with normal ACSF containing the following (in mM): 126 NaCl, 26 NaHCO_3_, 2.5 KCl, 1.25 NaH_2_PO_4_, 2 CaCl_2_, 2 MgCl_2_, 10 glucose (bubbled with 95% O_2_ and 5% CO_2_, pH 7.3; 310 mOsm measured). Lamina V putative WDR neurons were recorded based on the localization and the visualization of their large body size. Patch-clamp recordings were obtained with an Axon MultiClamp 200B amplifier coupled to a Digidata 1322A Digitizer (Molecular Devices, CA, USA). Borosilicate patch pipettes (2–4 MΩ) were filled with (composition in mM): 125 KCl, 10 HEPES, 2 MgCl_2_, 2 MgATP, 0.2 MgGTP (pH 7.3). ASIC currents were induced by locally puffing a MES-buffered pH6.6 solution for 5 seconds.

### Drugs

Synthetic mambalgin-1 was purchased from Synprosis/Provepep (Fuveau, France) and Smartox (Saint Martin d’Hères, France). PcTx1, MitTx and IbTx were purchased from Smartox. Apamin was purified in the laboratory from bee venom. Naloxone and morphine were purchased from Sigma-Aldrich (Saint Louis, MO, USA) and Copper (France), respectively. Toxins were prepared as stock solutions in ACSF, stored at -20°C, and diluted to the final concentration just before the experiments.

### Statistical analysis

Data are presented as the mean +/- SEM and statistical analysis was performed using Prism software. The statistical tests used to compare different sets of data are indicated in each figure legend.

### WDR neuron mathematical modeling including ASIC1a channel parameters and synaptic cleft acidification

Our computational model of WDR neuron (Fig.3 and Supplementary Figs 2-3) is an elaboration of a model introduced by Aguiar and colleagues (41). We refer to this work for the model description and only underline our modifications. The model, implemented in NEURON software, was downloaded from ModelDB (https://senselab.med.yale.edu/modeldb/).

By lack of evidence on the polysynaptic nature of the connection between the C-fiber and the WDR neuron, we removed the interneuron from the model in (41) to fit better the experimental spiking dynamics (see Results and Supplementary Fig. 2C-D). Our model is thus made of a WDR neuron, with dendrite, soma and axon, which receives a direct input from C-fiber via 20 synapses and from Aδ-fiber via 20 synapses. The C-fiber synapses each include AMPA, NMDA and GABA_A_ receptors with unchanged time dynamics and respective maximum conductances of 6nS, 4nS and 0.3nS, respectively, as well as a NK1 receptor with rise time constant of intermediate value *τ*_*rise*_ = 150*ms* and maximum conductance of 3pS. In order to finely adjust windup in the model without interneuron (before introducing ASICs) to experimental data, the calcium-dependent potassium current in the WDR model were tuned to 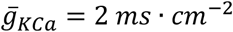 in the soma and 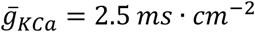 in the dendrite.

Next, we supplemented our WDR model with a model of ASIC1a channel, including ASIC1a homomers or heteromers (according to the most complete analysis of native ASIC channel parameters already done in spinal neurons (19)). The model is a simple Hodgkin–Huxley model of ionic channel, adapted from (42). The ASIC1a homomeric model is unchanged from the original publication, and the heteromeric/native model was obtained by modifying some of the parameters. As for the WDR model from (41), we refer to the corresponding publication for the structural and parametric choices that are not explicitly mentioned here. Note however that the null pH shift of 0.15 introduced in their Figure 2C was removed in both the homomeric and heteromeric ASIC1a models. In these models, the current flowing through the channels is defined as 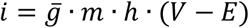, where *g* is the conductance of the channel, 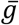 the maximal conductance, and *m* and *h* the activation and inactivation variables, respectively. Both variables are only sensitive to pH.

The heteromeric channel model was built to match the experimental data of “Type 2” native ASIC current from (19). The parameters for the asymptotic, pH-dependent values for *m* and *h* were modified to fit the values found in (19):

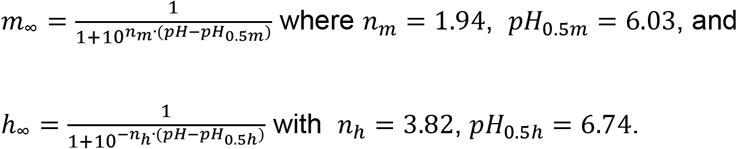

The order of magnitude of the time constant for the activation variable *m* is small enough to be considered instantaneous in our experimental setup, and its value as a function of pH was left unchanged from the ASIC1a model. The time constant τ_*h*_ of the inactivation variable *h* was fitted to data of (19). The rates for the relevant pH range (6.6-7.4) were plotted on a logarithmic scale and were found to be roughly aligned. Therefore a linear regression was performed on the log rates and the resulting expression of τ_*h*_ was:

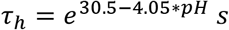

As done in (41) for NMDA and NK1 receptors, we modeled the ASIC1a-dependent increase in intracellular calcium concentration (50, 51) by setting 10% of the current entering through ASIC channels (both homomeric and heteromeric) as calcium current. The maximal ASIC conductance at each synapse varied between 0nS and 2nS for the purpose of our experiments, thus remaining always far below and at most half that of NMDA receptors.

Based on evidence of the postsynaptic involvement of ASICs in central synaptic transmission (44, 45), and of synaptic cleft acidification during transmission (52), we located ASIC channels in the WDR membrane at each synapse with the C-fiber. We modeled synaptic cleft acidification based on the work of Highstein and colleagues (43), with parameters fitted to our needs. The total proton concentration in the synaptic cleft is modeled as:

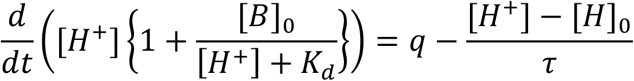

At every synaptic activation, defined by the presynaptic membrane potential going over -30mV, an input proton current q of 1ms duration is delivered into the cleft, modeling the putative proton sources during synaptic transmission (release of acidic vesicles, proton extrusion by the Na^+^/H^+^ exchanger…). The baseline proton concentration is given by the physiological pH as [*H*]_0_ = 10^−7.4^ *M*. The buffer parameters [*B*]_0_ = 22*mM* and *K*_*d*_ = 10^−6.3^ *M* were chosen for a model of physiological buffer used by (53). The values of the proton current q=0.5mM/ms and time constant τ = 0.5*ms*, least constrained by experimental data or physiological plausibility, were chosen from a range of acceptable values to adjust the resulting pH range to physiological values and to significantly affect windup (see supplementary figure 3). Using a similar grid search with the homomeric (rather than heteromeric) channel model, no values for q and τ could be found such that more than 20 spikes would be achieved on the last stimulation, as obtained with the experimental setup, unless the channel conductance was raised well above 1nS. Thus, the ASIC1a homomeric model was deemed unable to reproduce experimental results, unlike the heteromeric model which was therefore used for our experiments.

All simulation protocols were run with 15 stimulations delivered 1s apart starting at 1s, and the distribution of synaptic delays was kept unchanged.

The code for our WDR model, including ASIC channel parameters and the synaptic cleft acidification system, is available on request, and it will be uploaded to modelDB upon publication of this paper.

## Acknowledgements

We thank Drs A. Baron, S. Diochot, J. Noel, and M. Salinas for helpful discussions, V. Friend and J. Salvi-Leyral for technical support, and V. Berthieux for secretarial assistance. This work was supported by the Centre National de la Recherche Scientifique, the Institut National de la Santé et de la Recherche Médicale, the Association Française contre les Myopathies (AFM grant #19618), the Agence Nationale de la Recherche (ANR-11-LABX-0015-01, ANR-13-BSV4-0009 and ANR-17-CE16-0018) and the NeuroMod Institute of University Cote d’Azur (UCA).

## Author contributions

M.C. and L.P. designed, performed and analyzed *in vivo* spinal cord recordings. P.E. and K.D. designed, performed and analyzed patch-clamp experiments. A.D. and R.V. designed and realized the computational part of the work. E.L. helped in scientific design and critical reading of the manuscript. E.D. conceived the project, did data analysis and wrote the paper with the input of all authors.

## Competing interests

All authors declare no competing interests.

## Notes

### Competing Interest Statement

The authors have declared no competing interest.

