## supplementary material for "Involvement of ASIC1a channels in the spinal processing of pain information by deep projection neurons"

### SUPPLEMETARY FIGURES AND LEGENDS

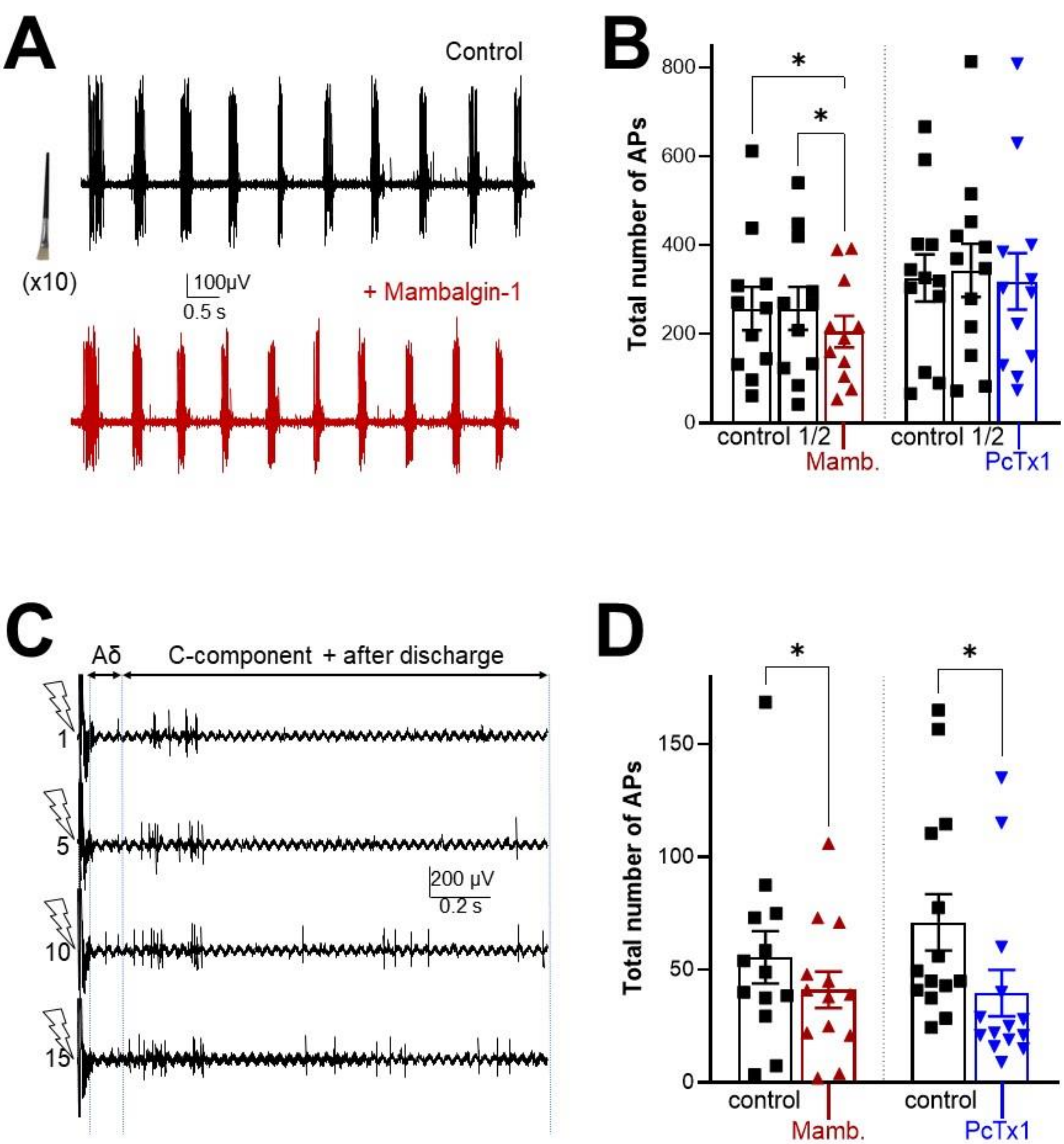


**<u>Supplementary figure 1</u>: Effect of spinal application of mambalgin-1 and PcTx1 on WDR neuron activities evoked by Aβ and δ fibers.** ***A,*** Typical recordings obtained following stimulation of a WDR neuron receptive field by 10 repetitive brushings (non-noxious stimulations), before (control) and after a 10-min application of mambalgin-1 (30µM) at the spinal cord level. ***B,*** The total number of AP evoked during the brushing protocol described in A are compared before (control 1 and 2) and after applications of either mambalgin-1 or PcTx1 30µM (n=13, *p*<0.05, one-way ANOVA tests followed by Dunnet's mutiple comparison tests).

***C,*** Typical recordings showing the activity of a WDR neuron during a windup protocol (16 repetitive electrical stimulations at 1Hz). Only the recordings obtained at stimulation 1, 5, 10 and 15 are represented. The vertical dashed lines represent the time ranges where the activity of the WDR is considered to be evoked by A$\delta$ or C fibres. ***D,*** Total number of AP evoked by A$\delta$ during windup protocols before (control) after applications of either mambalgin-1 or PcTx1 (n=13, *, $p$<0.01, paired t test).

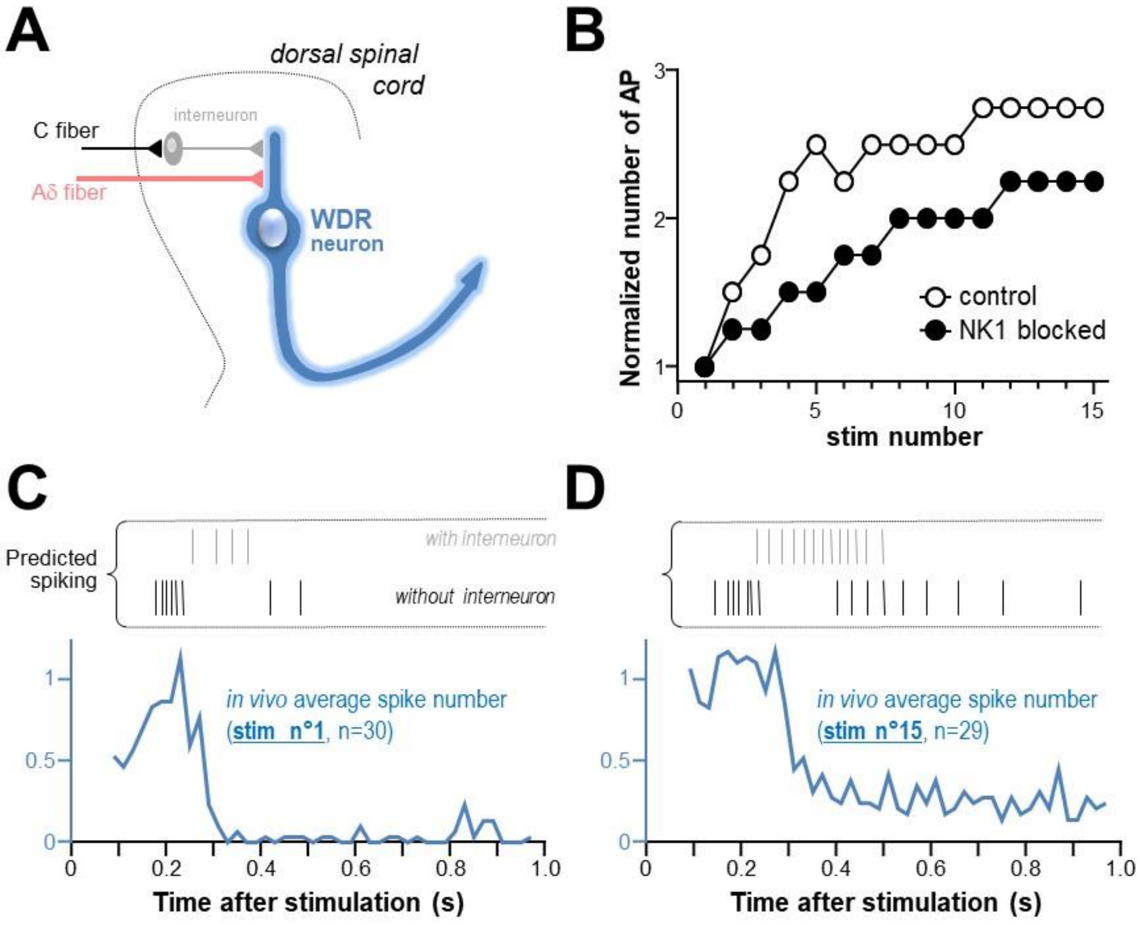


**<u>Supplementary figure 2</u>: Adapting the WDR neuron computational model.** ***A,*** Initial model developed by Aguiar and colleagues (41). ***B,*** Using Aguiar's model allows us to reproduce the results on windup when NK1 receptor parameters are removed. ***C-D,*** Spiking time profiles obtained for the first (stim1) and the fifteenth (stim 15) of the windup protocol (repetitive stimulations at 1Hz). Experimental data (blues curves), representing the mean number of AP per 20ms as a function of time (data from 29-30 neurons), are compared to data predicted by the Aguiar model, with (upper gray marks) or without (upper black marks) the interneuron between C-fiber and WDR neuron.

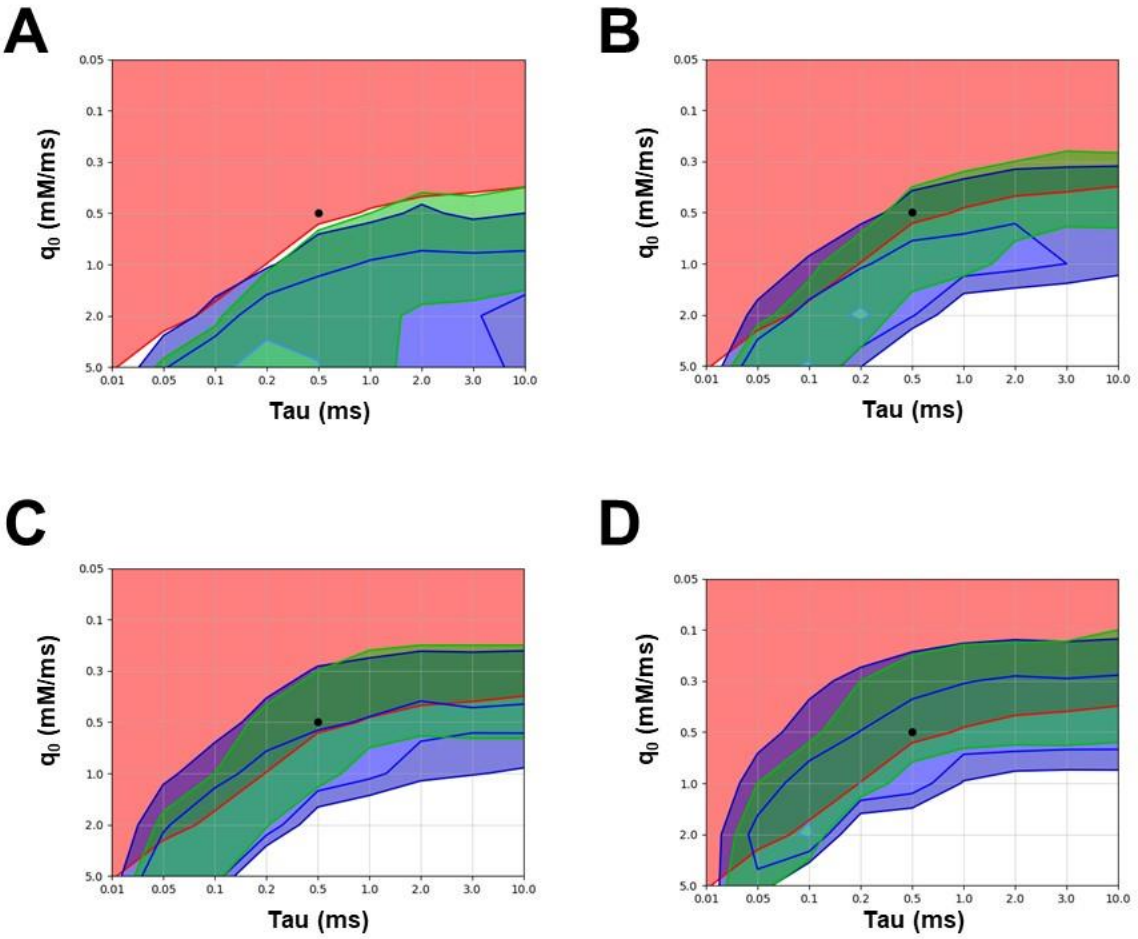


**<u>Supplementary figure 3</u>: Development of the pH model**

Grid searches for the values of q and $\tau$. The time constant $\tau$ has no direct physiological meaning and is unconstrained. The proton current q is considered acceptable below the maximum 2mM/ms, which represents the effect of an outward current of 250pA/pF (as was observed in HV1-transfected HEK293 cells depolarized to +90mV in (53)) through a membrane of capacitance 2,4µF/$cm^2$ as measured for spinal cord neurons by (54), into a synaptic cleft of 20nm width. We represent, on a q *vs.* $\tau$ plane, the zones in which *(i)* pH remains within a physiologically plausible range ($\geq$6.6, red zone), *(ii)* windup matches experimental data (blue zones, with threshold for AUC=142, 186 and 243) and (iii) the number of spikes elicited by the last stimulation matches experimental data ($\geq$22), this together with the previous condition constrains the shape of the windup curve, green zone. Black dot represents our chosen values. We made sure that the pH staying within the physiological range (with a small margin, pH≥6.5) for our chosen parameters was due to the parameters, rather than the interruption of stimulations before pH dropped further. This was tested by running a simulation with 100 stimulations and observing the stabilization of the pH minima.  ASIC maximal conductances are 0.1nS (***A***), 0.2nS (***B***), 0.3nS (***C***) and 0.5nS (***D***). For 0.2nS, the band of acceptable values is narrow; we chose a point in that band for our model (black dot). The green and blue zones move up with the conductance, resulting in a larger “acceptable” band. This suggests some physiological robustness to variations in ASIC quantity and synaptic cleft pH changes.
